# SARS-CoV-2 nsp3-4 suffice to form a pore shaping replication organelles

**DOI:** 10.1101/2022.10.21.513196

**Authors:** Liv Zimmermann, Xiaohan Zhao, Jana Makroczyova, Moritz Wachsmuth-Melm, Vibhu Prasad, Ralf Bartenschlager, Petr Chlanda

## Abstract

Coronavirus replication is associated with the remodeling of cellular membranes resulting in the formation of double-membrane vesicles (DMVs). Recently, a pore spanning DMV was identified as a putative portal for viral RNA transcription and replication products providing a novel target for antiviral intervention. However, the exact components and the structure of the SARS-CoV-2 pore remain to be determined. Here, we investigate the structure of DMV pores by *in situ* cryo-electron tomography combined with subtomogram averaging. We reveal non-structural proteins (nsp) 3 and 4 as minimal components forming a DMV spanning pore and show that nsp3 Ubl1-Ubl2 domains are critical for inducing membrane curvature and DMV formation. Altogether, SARS-CoV-2 nsp3-4 has a dual role by driving the biogenesis of replication organelles and forming DMV-spanning replicopores.

**One-Sentence Summary:** Biogenesis of SARS-CoV-2 replication organelles is driven by nsp3-4 constituting the double-membrane vesicle spanning pore.

## Main Text

Positive-strand RNA viruses hijack and remodel the host cell membranes into replication organelles gated by proteinaceous pores *(1–3)*. Severe acute respiratory syndrome coronavirus 2 (SARS-CoV-2) induces the formation of endoplasmic reticulum (ER)-derived double-membrane vesicles (DMVs), which serve as replication organelles (ROs) *(4–6)*. It is well established that coronavirus replication-transcription complex (RTC) is associated with DMVs to orchestrate viral genome replication and transcription yielding subgenomic messenger RNAs (sgRNAs) using double-stranded (ds) RNA as intermediate *(4, 6, 7)*. While dsRNA can be directly visualized inside the DMVs *(8)* and detected by immunolabelling *(4, 6)* it is not clear whether the replication and transcription of viral RNA take place on the inner (luminal) or outer (cytoplasmic) side of the DMV. DMVs may provide a shielded environment for viral RNA, evading recognition of innate immune sensors present in the host cell and thereby facilitating robust viral genome replication and transcription. Recently a structure spanning the DMV induced by the murine hepatitis coronavirus (MHV) was identified and proposed to serve as a putative pore dedicated to the translocation of the newly synthesized viral genomic RNA (gRNA) and sgRNAs from the DMV lumen into the cytoplasm *(1)*. However, the structure of the SARS-CoV-2 pore has not been determined yet and the exact constituents and minimally required protein components of the pore complex are still enigmatic. While the non-structural protein (nsp) 3 is the largest multi-domain protein encoded by the coronavirus genome and presumably the major component of the pore complex that constitutes the crown region of the pore as shown for MHV *(1)*, the architecture of the pore spanning DMVs in SARS-CoV-2 infected cells is unknown. It has been shown that the Middle East Respiratory Syndrome coronavirus (MERS-CoV), SARS-CoV and SARS-CoV-2 nsp3 and nsp4 are sufficient to induce DMV formation *(9–13)* and it remains to be determined whether the expression of nsp3 and nsp4 is sufficient to form a pore. Moreover, the mechanism of how these proteins induce membrane curvature to shape ER into DMVs and whether the assembly of the pore is concomitant with membrane bending remains to be elucidated. Here we established a workflow that allows us to investigate the structure of the SARS-CoV-2 DMV spanning pore and to determine its minimal components using cryo-electron tomography (cryo-ET) and subtomogram averaging in the native cellular environment and BSL1 conditions. Hence, circumventing the necessity of chemical inactivation applied on SARS-CoV-2 infected cells prior to plunge freezing for biosafety reasons.

To determine the minimally required components assembling a SARS-CoV-2 pore spanning DMVs, we performed *in situ* cryo-ET on lamellae prepared by focused ion beam (FIB) milling of transfected cells expressing SARS-CoV-2 nsp3-4. To increase our throughput, a plasmid encoding the fluorescent protein mApple was co-transfected with the nsp3-4 construct. Our previous work showed that co-transfection of different contracts leads to efficient co-expression of proteins in transfected cells *(14)*. This allowed us to perform cryo-correlative light and electron microscopy to target only transfected cells by cryo-FIB milling (Fig. S1). We observed a network of DMVs in the cytoplasm of VeroE6 cells clearly showing pores spanning the DMVs (Fig. 1A-B, Movie S1). 3D segmentations revealed that nsp3-4 induced DMVs are interconnected by double membrane sheets which we termed double membrane connectors (DMCs) (Fig. 1A-B, E, Fig. S2A). In addition, DMVs were frequently connected to ER cisternae decorated by ribosomes, indicating that the DMV network induced by nsp3-4 expression originates from ER membranes similarly as reported for DMVs induced upon SARS-CoV-2 infection *(15)* (Fig. S2B-C). The mean diameter of DMVs formed by nsp3 and nsp4 was 104 nm (SD = 50, *n* = 62) thus smaller than the DMV diameter of ∼336 nm (SD = 50, *n* = 20) measured in infected VeroE6 cells (Fig. S3B), consistent with previous reports *(12, 15)*. Compared to the DMVs found in infected cells, the lumen of the nsp3-4 induced DMVs did not contain filamentous material indicating that nsp3-4 alone is not able to transport cellular RNA into the DMV lumen. Moreover, vesicle packets formed by homotypic DMV-DMV membrane fusion reported previously during late infection *(4, 8)*, were not detected in nsp3-4 induced DMVs. Additionally, we observed that some DMVs contain densities (Fig. S4), which are similar to those described as dense granules inside DMVs found in poliovirus infected cells suggesting the involvement of autophagic membranes or proteins in DMV biogenesis *(16)*. Importantly, due to the lack of chemical fixation, the structure of the nsp3-4 pores was better preserved in transfected cells than in SARS-CoV-2 infected cells *(1, 8)* (Fig. 1C, Fig. S3A). Strikingly, both DMVs and DMCs contained clearly discernable pores which consist of a double membrane spanning region and of a crown-like assembly facing the convex side of the DMV (Fig. 1C, F). Subtomogram averaging revealed a pore structure at a resolution of 20 Å with sixfold rotational symmetry along the axis normal to the DMV surface (Fig. 1G-H, Fig. S5). The crown part of the pore showed a 25-nm-wide prong platform decorating the convex cytosolic side of the DMV. Underneath the prongs, a channel of 2-3 nm diameter was detected, which is the direct connection to the luminal side of the DMV (Fig. 1I). The overall architecture of the SARS-CoV-2 nsp3-4 pore shares structural similarity with pores in DMVs resulting from MHV infection *(1)* (Fig. S6D). This demonstrates that the expression of nsp3 and nsp4 is sufficient for the formation of the SARS-CoV-2 DMV pore.

**Fig. 1.**
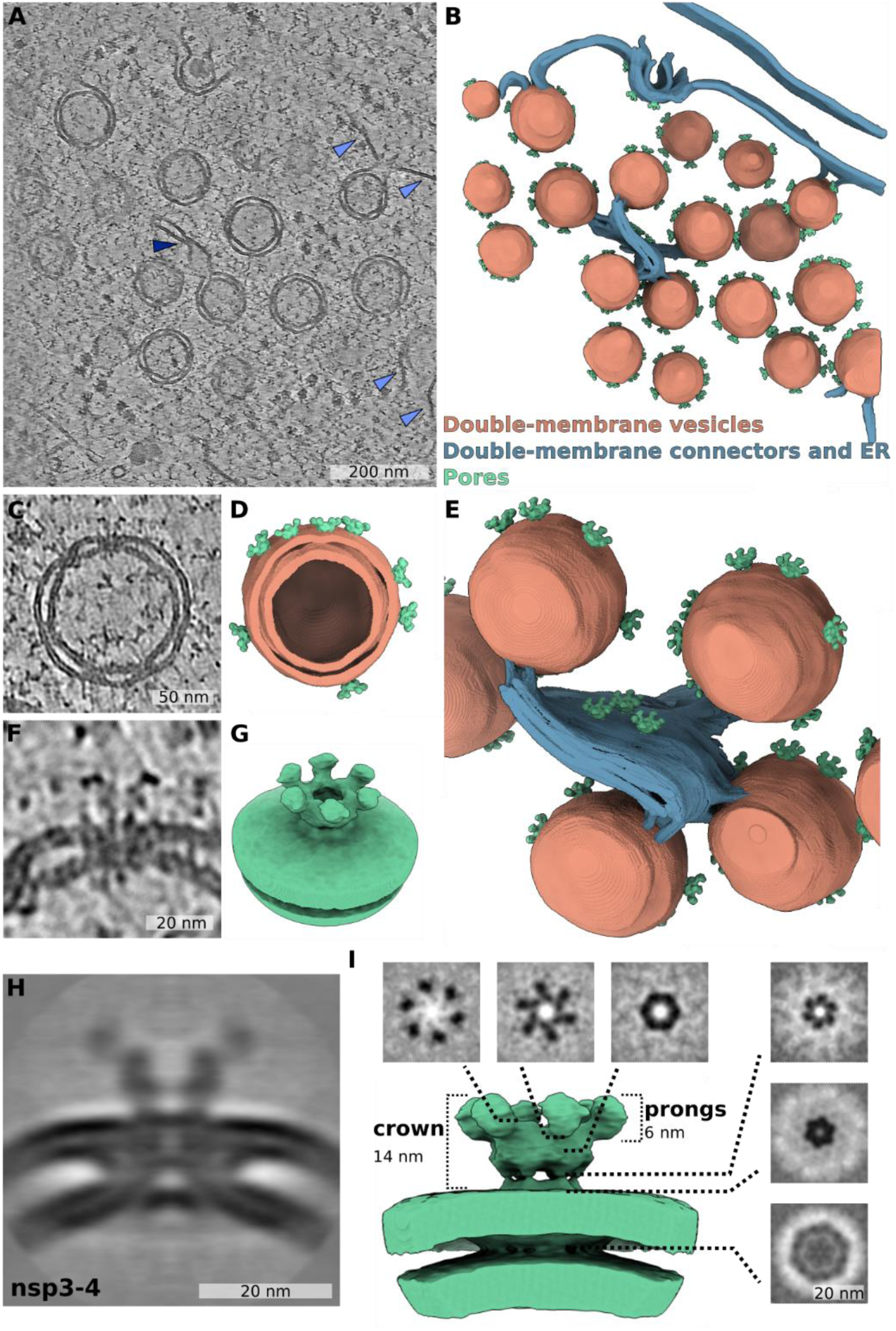
The SARS-CoV-2 nsp3-4 proteins are sufficient to induce DMV formation and DMV-spanning pores. (A) Averaged slices of a tomogram acquired on cryo-lamella of VeroE6 cells transfected with HA-nsp3-4-V5 and plunge frozen at 16 hpt. The double-membrane vesicles (DMVs) are interconnected through double-membrane connectors (DMCs) which are highlighted by dark blue arrows. The connections between DMVs and ribosome-decorated endoplasmic reticulum are highlighted by light blue arrows. (B) Volume rendering of DMV and DMC network with pores spanning the DMVs and the DMCs. (C) Averaged slices of a tomogram showing a zoomed area of a single DMV with multiple pores spanning two membranes in (A). (D) Volume rendering of the DMV in (C) showing the inside of the DMV. (E) Volume rendering showing a zoomed area of DMVs interconnected through pore containing DMCs in (B). (F) Averaged slices of a tomogram showing a zoomed in area of one DMV pore in (C). (G) Isosurface of the filtered C6 symmetrized subtomogram average of SARS-CoV-2 HA-nsp3-4-V5 showing the pore complex with its crown sitting on the convex side of the double membrane. (H) Slices averaged through the filtered C6 symmetrized subtomogram average of the pore complex. (I) Slices through the filtered C6 symmetrized subtomogram average from top to bottom. Isosurface of the pore complex induced by SARS-CoV-2 HA-nsp3-4-V5 (green).

To further investigate the structure of the pore and the role of nsp3 in DMV biogenesis, we generated two N-terminal nsp3 truncations by deleting (i) the ubiquitin-like domain 1 (Ubl1), hypervariable region (HVR) and macrodomain 1 (Mac1) (construct ΔUbl1-Mac1) or (ii) Ubl1, HVR, Mac1-3, Domain Preceding Ubl2 and PL2^pro^ (DPUP) and Ubl2 (construct ΔUbl1-Ubl2). In addition, to shed light on the importance of the proteolytic separation of nsp3 and nsp4 to the pore formation, we substituted GG for AA at the cleavage site at position 1944-1945 (GG>AA), which renders the polyprotein unable to cleave *(10)* (Fig 2A). Western blot analysis of all nsp3-4 constructs showed bands consistent with the predicted molecular masses and, except for the GG>AA construct, proteolytic processing at the nsp3-4 GG cleavage site mediated by PL2^pro^ (Fig. 2A-B). Interestingly, the deletion in the ΔUbl1-Ubl2 construct slightly reduced nsp3-4 cleavage, indicating that the Ubl2 domain proximal to PL2^pro^ is supporting the function of the protease or contributes to proper exposure of the cleavage site (Fig 2B). Confocal microscopy revealed that nsp3 and nsp4 colocalize in all constructs except the ΔUbl1-Ubl2 construct in which the nsp3 signal was weak, presumably because of impaired folding and only partial exposure of the HA-tag at the N-terminus (Fig. 2D-G). Expression of all nsp3-4 constructs led to membrane remodeling in cells as determined by electron microscopy analysis of samples processed by high pressure freezing and freeze substitution (Fig. 2H-K). Consistent with studies on MERS-CoV *(10)*, abrogation of SARS-CoV-2 nsp3-4 cleavage resulted in the formation of large double-membrane whorl-like structures while DMVs were not detected. This indicates that nsp3-4 proteolytical cleavage is followed by rearrangements of nsp3 and nsp4 providing the required membrane energy for the biogenesis of DMVs.

**Fig. 2.**
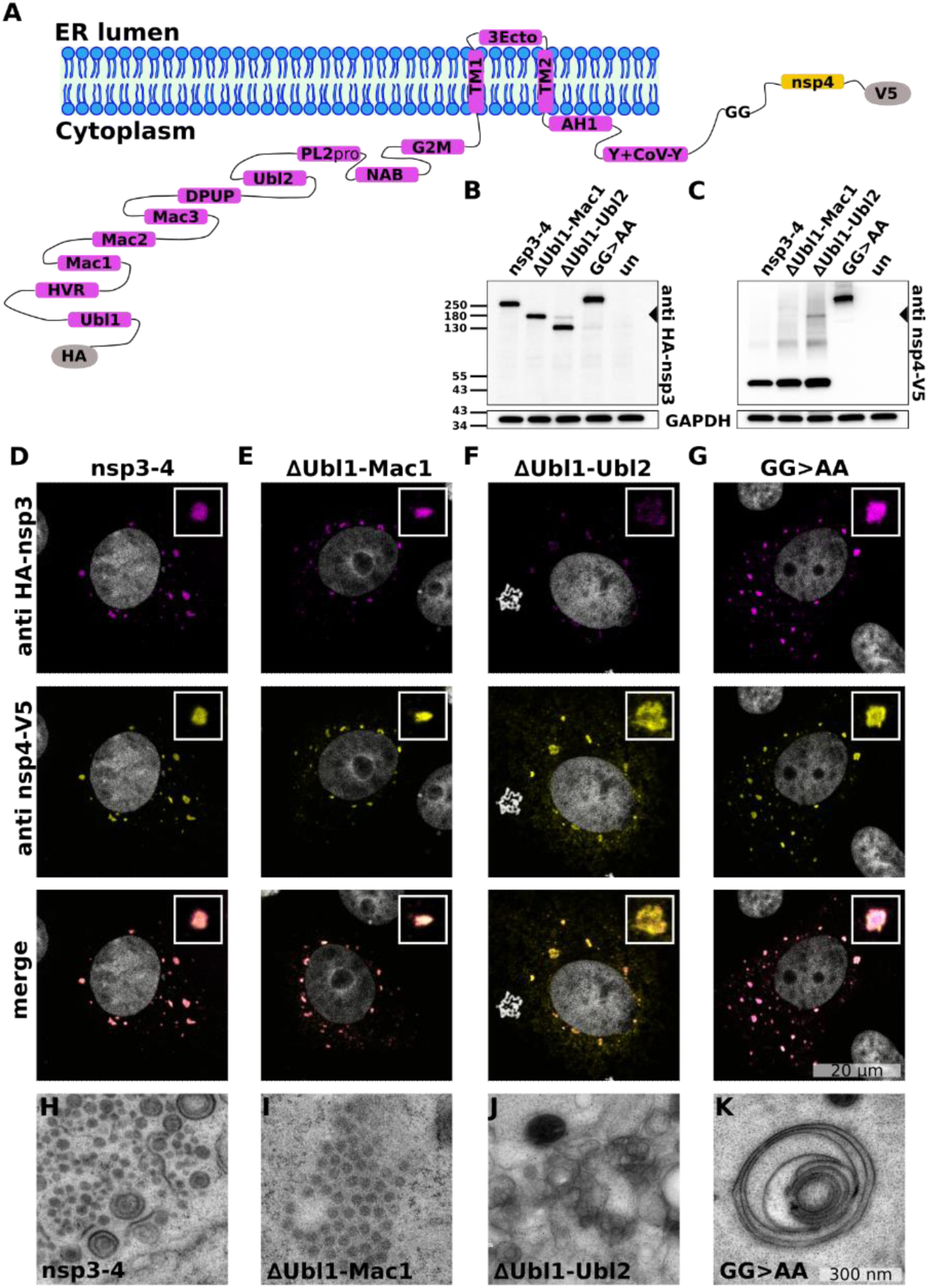
The N terminal domains of SARS-CoV-2 nsp3 are critical for the formation of distinct DMVs. (A) Schematic representation of membrane bilayer with nsp3-4 polyprotein tagged with hemagglutinin tag at the N-terminus and V5 tag at the C-terminus. Nsp3 domains and nsp4 are shown in magenta and in yellow, respectively. The cleavage site between nsp3 and nsp4 polyprotein is indicated by GG. Nsp3-4 truncations were generated by removal of N-terminal domains ΔUbl1-Mac1 and ΔUbl1-Ubl2 and the cleavage mutant GG>AA was generated by substituting GG with AA at the position 1944-1945. All constructs were tagged at the N- and C-termini, with HA and V5 tags, respectively. Western blot analysis, using anti-HA (B) and anti-V5 (C), of lysates of untransfected VeroE6 cells (un) and VeroE6 cells transfected with nsp3-4, ΔUbl1-Mac1, ΔUbl1-Ubl2, or GG>AA construct harvested at 24 hpt. Additional bands present in ΔUbl1-Ubl2 in C and B are indicated by an arrow. (D-G) Confocal microscopy of VeroE6 cells transfected with nsp3-4 (D), ΔUbl1-Mac1 (E), ΔUbl1-Ubl2 (F) and GG>AA (G) chemically fixed at 24 hpt (representative images of nsp3-4: n = 17; ΔUbl1-Mac1: n = 11; ΔUbl1-Ubl2: n = 13; GG>AA: n = 10 cells). Nsp3 and nsp4 were detected by indirect immunofluorescence using anti-HA and anti-V5 antibodies, shown in magenta and yellow, respectively. (H-K) Thin-section EM images of HEK293T cells transfected with respective constructs and high-pressure frozen at 24 hpt.

We next performed *in situ* cryo-ET on cryo-FIB milled lamellae of transfected VeroE6 cells to assess the membrane remodeling and determine whether nsp3-4 truncated at the N-terminus is sufficient for pore formation. Consistent with the data from thin-section EM, both nsp3-4 truncations were able to induce membrane remodeling. The expression of the ΔUbl1-Mac1 construct led to the formation of DMVs that contained a similar number of pores but had a larger DMV radius than DMVs induced by unaltered nsp3-4 (Fig. 3A-B, E-F, Fig S6E-F, Movie S2). In contrast, the ΔUbl1-Ubl2 nsp3-4 truncation was not able to remodel membranes into distinct DMVs, but instead, we observed a double-membrane network composed of DMCs, which contained pores and formed only partially closed ovoidal structures (Fig. 3C, G, Movie S3). Cryo-ET data allowed us to estimate the number of pores per DMV, and measure pore-to-pore distance and DMV luminal spacing, which is the distance between the two membranes forming DMVs (Fig. 3I-L). DMVs formed by nsp3-4 contained on average 11 (SD = 2, *n* = 20) pores per DMV, which is approximately two times higher than the number of reported pores per DMVs formed in MHV infected cells (∼5 pores per DMV) *(1)*. Considering that DMVs formed in infected cells are larger, a higher number of pores per DMV may be directly proportional to increased membrane bending and thus smaller DMVs. Interestingly, the double membrane whorl-like structures induced by the cleavage incompetent mutant of nsp3-4 (GG>AA) did not contain any pores (Fig. 3D, H, Movie S4) and the luminal spacing was reduced from 16 nm (SD = 1.1, *n* = 18) to 13 nm (SD = 0.8, *n* = 19). This indicates that the uncleaved nsp3-4 induces membrane stacking, but pore formation is required for membrane bending and scission which gives rise to DMV.

**Fig. 3.**
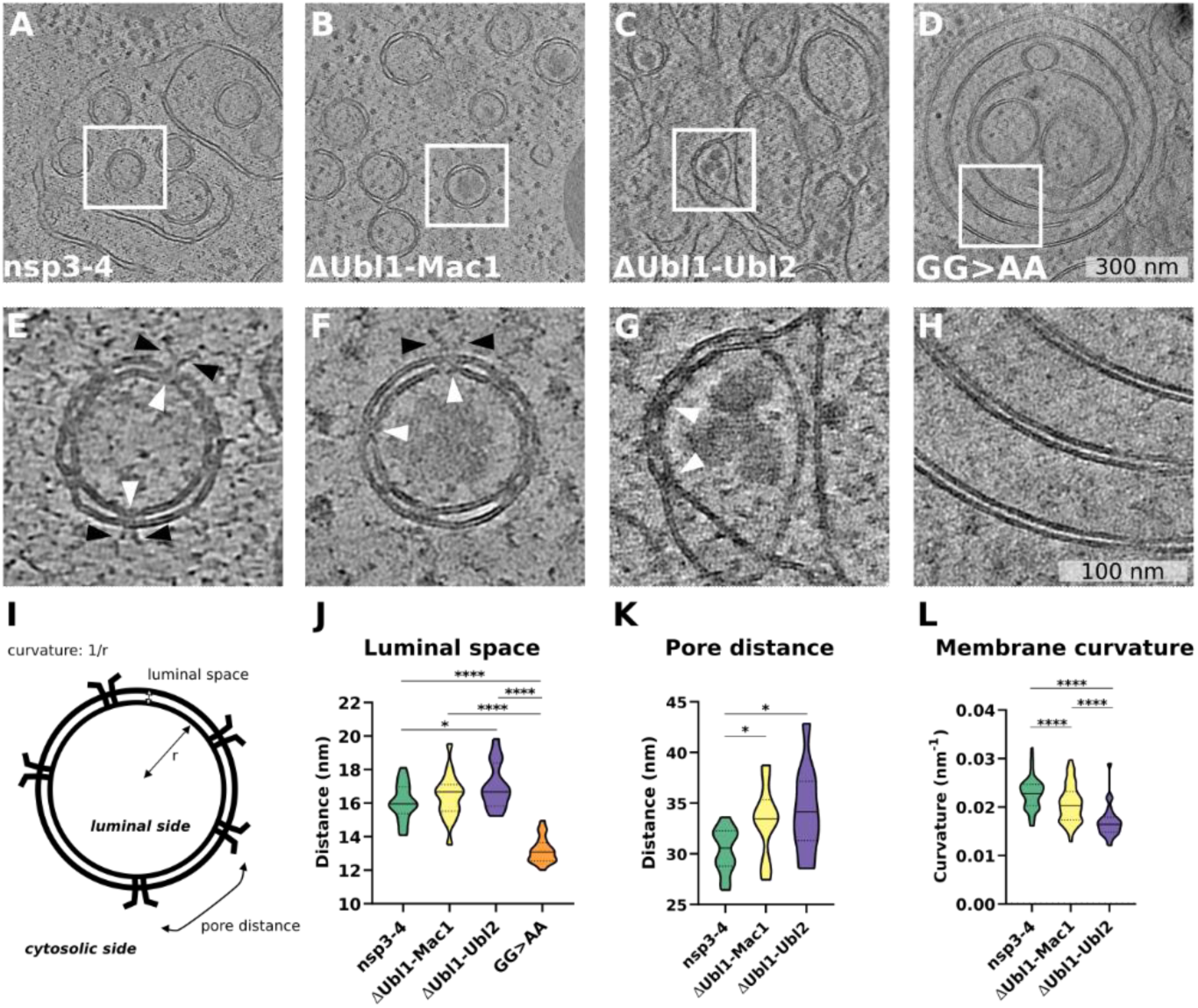
Domains at the SARS-CoV-2 nsp3 N-terminal region are critical for the formation of DMV-spanning pores. (A-D) Slices of cryo-electron tomograms of VeroE6 cells transfected with nsp3-4 (A), ΔUbl1-Mac1 (B), ΔUbl1-Ubl2 (C) and GG>AA (D) plunge frozen at 18-24 hpt. (E-H) Slices of tomograms displaying magnified areas in A-D. White arrowheads indicate pores spanning the DMVs, prongs are indicated by black arrowheads. (I) Schematic representation of DMV and indicated parameters that are displayed in plots shown in (J-L). (J) Luminal spacing between two membranes of a DMV (nsp3-4: n = 18; ΔUbl1-Mac1: n = 17; ΔUbl1-Ubl2: n = 15; GG>AA: n = 19). (K) Pore-to-pore distance on the surface of DMV (nsp3-4: n = 12; ΔUbl1-Mac1: n = 12; ΔUbl1-Ubl2: n = 11). (L) Membrane curvature of DMV determined using osculating circle method (nsp3-4: n = 103; ΔUbl1-Mac1: n = 103; ΔUbl1-Ubl2: n = 26). Data is shown as Violin plot indicating the median, 25% and 75% quartiles. Unpaired two-tailed t-test showed significant differences with *p<0.03, ****p<0.0001.

To compare the architecture of DMV pores formed by truncated nsp3-4 constructs, we applied subtomogram averaging of pores detected with ΔUbl1-Mac1 and ΔUbl1-Ubl2 induced DMVs, respectively. Our data showed that while pores formed by nsp3-4 lacking N-terminal Ubl1-Mac1 domains still formed crown-like structures on the convex side of the DMVs, the prongs were partially collapsed and shorter at the distal end in comparison to wildtype nsp3-4 pores, presumably because of altered inter-nsp3 interactions leading to instability of the prongs (Fig. 4A, B, D, E, Fig. S6A). Surprisingly, nsp3-4 lacking the Ubl1-Ubl2 domain showed no crown-like structure and only a membrane-proximal density of prongs was visible (Fig. 4C, F, Fig. S6B). This data demonstrates that the nsp3 N-terminal region including domains localized between Ubl1 and Ubl2 is critical for the formation of the crown-like structure and pivotal for nsp3-nsp3 interactions, which are likely required for multimerization. Subtomogram averaging highlighted the difference in membrane curvature formed by different nsp3-4 constructs. While nsp3-4 was able to induce membrane curvature J=0.017 nm^-1^ of the outer membrane (Fig. 4G), truncations of the nsp3 N-terminus, which are associated with loss of crown structural integrity, caused flattening of the outer membrane. Strikingly, the deletion spanning Ubl1-Ubl2 domains led to a reduction of membrane curvature to J=0.008 nm^-1^. Overall, our data show that SARS-CoV-2 nsp3-4 is sufficient to form a pore that spans DMVs. While the pore shares global structural similarities to the pore found in MHV infected cells *(1)*, the structure of the prongs is different in the SARS-CoV-2 nsp3-4 transfected cells compared to the prongs found in MHV infected cells (Fig. S6D). The results support the notion that coronavirus pores are similar in structure and are composed of nsp3-4. However, the N-terminal domain organization of MHV nsp3 is different when compared to the N-terminal domains of SARS-CoV-2 nsp3. MHV nsp3 does not contain the Mac2 and Mac3 domains which might explain the structural differences in the prongs *(17)*. The structural data on nsp3 truncations allow us to propose a model on stepwise biogenesis of DMVs where nsp3 and nsp4 have distinct but synergistic roles. On the one hand, nsp4 is critical in establishing a double membrane sheet composed of two parallel membranes and is required to induce strong local membrane bending leading to a pore in resemblance to the assembly of nucleopore complexes *(18)*. On the other hand, the architecture of the nsp3 N-terminal crown contributes to the bending of the membrane sheets leading to the formation of distinct DMVs. It remains to be seen, whether the deletion of the nsp3 crown structure leads to conformational changes or rearrangement of nsp4, which in turn presumably controls membrane curvature. In addition, nsp3 engages in sensing and recruitment of viral RNA *(19)* in analogy to the nucleopore cytoplasmic fibrils selecting protein cargo for import. In analogy to nucleopores, we propose to term the proteinaceous pores induced by coronaviruses and other positive-strand RNA viruses “replicopores”. Finally, the established experimental framework for reconstitution of SARS-CoV-2 replication *in situ* will be instrumental to investigate other putative structural components of the replicopore such as RTC and opens an avenue for high-resolution determination.

**Fig. 4.**
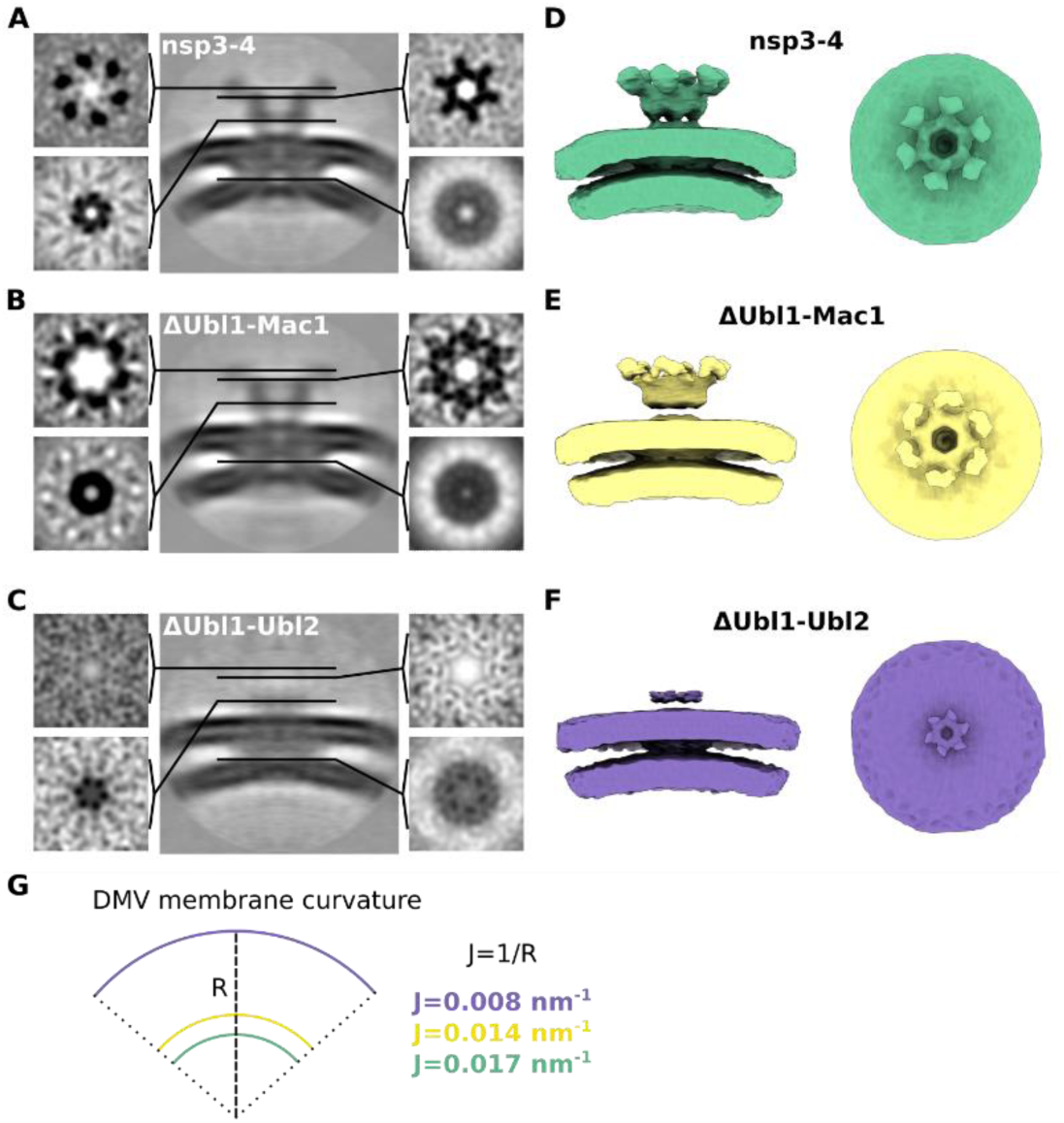
The region spanning Ubl1-Ubl2 domains of SARS-CoV-2 nsp3 contribute to membrane curvature. (A-C) Orthogonal slices of filtered C6-symmetrized subtomogram averages obtained from cryo-electron tomograms of VeroE6 cells transfected with nsp3-4 (A), ΔUbl1-Mac1 (B), ΔUbl1-Ubl2 (C). The position of orthogonal slices along the z axis is indicated by black lines in the cross-sectional view of the average. (D-F) Side (left) and top views (right) of filtered subtomogram averages displayed as isosurfaces. (G) Membrane curvature of DMV outer membrane changes upon truncation of nsp3.

## Acknowledgments

We thank the Infectious Diseases Imaging Platform (IDIP) at the Center for Integrative Infectious Disease Research Heidelberg and the cryo-EM network at the Heidelberg University (HD-cryoNET) for support and assistance. The authors gratefully acknowledge the data storage service SDS@hd supported by the Ministry of Science, Research, and the Arts Baden-Württemberg (MWK), the German Research Foundation (DFG) through grant INST 35/1314-1 FUGG and INST 35/1503-1 FUGG.

## Author contributions

Conceptualization: LZ, XZ, PC; Methodology: LZ, XZ, JM, MWM, VP, PC; Investigation: LZ, XZ, JM, PC; Visualization: LZ, XZ, JM; Funding acquisition: RB, PC; Project administration: JM, PC; Supervision: RB, PC; Writing – original draft: LZ, XZ, PC; Writing – review & editing: LZ, XZ, JM, MWM, VP, RB, PC

## Data and materials availability

Electron tomography data have been deposited to the Electron Microscopy Data Bank under accession codes EMD-15925 (tomogram of nsp3-4 induced DMVs in Fig.1), EMD-15926 (tomogram of ΔUbl1-Ubl2 induced pores in Fig.3), EMD-15927 (tomogram of ΔUbl1-Mac1 induced pores in Fig.3), EMD-15928 (tomogram of GG>AA induced pores in Fig.3), EMD-15929 (tomogram of nsp3-4 induced pores in Fig.3), EMD-15963 (subtomogram average of nsp3-4 pore in Fig.1), EMD-15964 (subtomogram average of ΔUbl1-Mac1 pore in Fig.4), EMD-15965 (subtomogram average of ΔUbl1-Ubl2 pore in Fig.4) and will be available upon publication. Additional data and material related to this publication may be obtained upon request.

