## Supplementary Materials for "SARS-CoV-2 nsp3-4 suffice to form a pore shaping replication organelles"

### Materials and Methods:

#### Cell culture

HEK293T (293T ECACC, 1202201) cells were purchased from Sigma-Aldrich. VeroE6 cells were purchased from American Type Culture Collection (ATCC; Catalogue #CRL-1586). Cells were maintained in Dulbecco's modified Eagle medium (DMEM, Thermo Fisher Science) with high concentration of glucose, GlutaMAX supplement, 100 U/ml penicillin, 100 µg/ml streptomycin, and 10% fetal bovine serum at 37°C and 5% $CO_2$. For DNA transfection TransIT-LT1 Transfection Reagent was used according to the manufacturer's protocol (Mirus Bio LLC).

#### Plasmids and antibodies

Codon-optimized synthetic DNA of 7416 bp expressing HA-nsp3-nsp4-V5 cDNA was ordered in 3 fragments from Biocat GmbH (Heidelberg) and assembled using overlap-extension PCR followed by insertion in pcDNA3.1 vector backbone using EcoRI-XbaI restriction sites. The expression of nsp3 and nsp4 from this plasmid and the induction of DMVs were confirmed previously (*20*). Truncations were generated using In-Fusion HD Cloning Plus kit (TAKARA Bio). The sequences of the nsp3-4 constructs used in this study are listed in Supplementary Data S1. pN1-mApple plasmid (#54567) was purchased from Addgene. To detect the HA-tag and V5-tag by confocal microscopy and Western blot analysis, a rabbit anti-HA (Invitrogen, 71-5500) and mouse anti-V5 antibodies (Santa Cruz Biotechnology, sc-271944) were used as primary antibodies, respectively. The secondary antibodies Alexa Fluor 546 goat anti-rabbit (Invitrogen, A11010) and Alexa Fluor 488 goat anti-mouse (Invitrogen, A11029) were used for confocal microscopy. The secondary antibodies mouse anti-rabbit IgG-HRP (Santa Cruz Biotechnology, sc-2357) and anti-mouse IgG BP-HRP (Santa Cruz Biotechnology, sc-516102) were used for Western blot analysis.

### Western blot analysis

VeroE6 cells were seeded into 6-well plates at a seeding density of 0.4 × $10^6$ cells/well. Cells were transfected with 2µg DNA/well. At 24 hours post transfection (hpt), cells were washed with cold PBS and lysed using 1 ml prechilled lysis buffer (2% SDS, 50 mM Tris-HCl, pH 7.4) supplemented with protease inhibitor cocktail in PBS (Roche) for 15 min. The cell lysate was centrifuged at 16,000 × g, 15 min, 4°C. The supernatant was collected and mixed with Laemmli buffer (BioRad) and DTT (final concentration 50 mM). All samples were boiled at 95 °C for 10 min and separated by electrophoresis on SDS-polyacrylamide gradient gels (4-15%). Proteins were transferred to polyvinylidene fluoride (PVDF) membranes (Bio-Rad) using a Trans-Blot Turbo transfer system (Bio-Rad). Membranes were blocked with 5% milk in TBS-T (1 × TBS with 0.1% Tween-20) for 1 h at room temperature (RT). The PVDF membrane was washed for 3 ×10 min with TBS-T and incubated in primary antibody solutions prepared by diluting 1:1000 in TBS-T with 5% milk for 1 h at RT. Subsequently, membranes were washed 3 times with TBS-T and incubated in secondary antibody solutions prepared by diluting 1:1000 in TBS-T with 5% milk for 1 h at RT. Blots were washed 3 × 10 min with TBS-T and incubated in enhanced chemiluminescence substrate (Clarity Western ECL substrate, Bio-Rad) solution for 5 min at RT in dark. Images were acquired with Azure 400 Imaging System (Azure Biosystems).

### Confocal microscopy

VeroE6 cells were seeded into 12-well plates on coverslips at a seeding density of 0.12 × $10^6$ cells/well. Cells were transfected with 750 ng DNA/well. Cells were washed with PBS at 24 hpt and fixed by 4% formaldehyde (PFA) diluted in PBS (16% PFA (E15710), Science Services) for 15-30 min at RT. Cells were washed twice with PBS and permeabilized with permeabilization buffer (0.5% Triton X-100 in PBS) for 5 min at RT. Cells were washed 3 × 5 min with PBS and blocked with blocking buffer (3% lipid-free BSA in PBS-T) for 1 h at RT. Cells were washed 3 times with PBS and incubated with primary antibody solutions prepared by diluting 1:200 in dilution buffer (1% lipid-free BSA in PBS-T) for 1 h at RT. Cells were washed 3 × 5 min with PBS and incubated in secondary antibody solutions prepared by diluting 1:500 in dilution buffer for 1 h at RT in dark. Cells were washed 3 × 5 min with PBS and incubated in DAPI mix (1:1000 dilution in PBS) for 1 min at RT. Cells were washed 2 × 5 min with PBS and once with deionized water and mounted on clean glass slides using a 7 µl mounting medium (Prolong

Glass, Life Technologies). After drying, samples were analyzed with Leica TCS SP8 confocal laser scanning microscope, equipped with a 63× objective (numerical aperture 1.40; 1 Airy unit) and a Leica HyD hybrid detector. One slice was chosen to show in the images, and the brightness and contrast were adjusted to the same scale in ImageJ/Fiji (*21*).

**High pressure freezing and freeze substitution**

Sapphire discs with 3.0 mm diameter and 50 μm thickness (Wohlwend GmbH) were cleaned with 100% ethanol and coated with a 15 nm carbon layer using a sputter coater (EM ACE600, Leica). An "F" letter was scratched on the carbon side to distinguish between the two sides of the disc, followed by overnight incubation of the discs at 120°C. Prior to use, the carbon-coated discs were plasma-cleaned for 10 seconds (s) in a Solarus 950 (Gatan), sterilized in 70% ethanol, washed once with DMEM, and placed in 1 ml DMEM in a 35 mm dish with the "F" side facing up. The DMEM was removed and HEK293T cells were seeded on the sapphire discs ($0.18 \times 10^6$ cells in 2 ml DMEM per dish). Cells were transfected with 1 µg DNA/dish (co-tranfection of pN1-mApple plasmid). At 24 hpt, sapphire discs with transfected cells were assembled with 1-hexadecene coated specimen carrier Type A and B (Wohlwend GmbH) with cells facing the 100 µm deep cavity of carrier Type A. Cells were vitrified by high-pressure freezing at approximately 2,200 bar maintained for 370 ms with a cooling rate of 20,000 K/s using a Leica EM ICE. Sapphire discs with vitrified cells were transferred from liquid nitrogen to a freeze-substitution (FS) solution (0.1% uranyl acetate in anhydrous acetone) cooled to -90°C and processed in automated FS system (EM AFS2, Leica). After washing with acetone, samples were infiltrated with Lowicryl HM20 and polymerized using UV light. The FS protocol was performed according to Table 1.

**Ultramicrotomy and electron microscopy of resin sections**

Lowicryl-embedded samples were sectioned using diamond knives (DiATOME) and a UC7 ultramicrotome (Leica). Sections with 200 nm nominal thickness were placed on 2 × 1 mm copper slot grids (Gilder) coated with support film (1% formvar or pioloform). Grids were imaged with a Talos L120C TEM operated at 120 keV and equipped with a Ceta-M camera with a 4k × 4k CMOS (Thermo Fisher Scientific). The whole grid was mapped at a magnification of 155 × and images were acquired at magnifications of 5,300×; 11,000× and 45,000×

(corresponding pixel sizes at the specimen level: 26.44 Å, 13.35 Å and 3.28 Å respectively) using SerialEM (*22*).

**Plunge-freezing**

To prepare samples for plunge-freezing, 35 mm cell culture dishes coated with a thin layer of polydimethylsiloxane (PDMS) were used to culture cells. Holey carbon grids (200 mesh Quantifoil™ Au R2/2 grids) were placed into a PDMS coated dish and plasma-cleaned for 10 s in a Gatan Solarus 950 (Gatan). The PDMS-coated dish with grids was sterilized with 70% ethanol and washed twice with DMEM. The DMEM was removed and VeroE6 cells were seeded on the grids at a seeding density of $0.12 \times 10^6$ cells/dish. Cells were transfected with 1 µg DNA/dish (925 ng of nsp3-4 constructs, 75 ng of mApple construct). At 18-24 hpt, cells were stained with 1 µg/ml Hoechst (B2261) for 5 min and washed once with DMEM. Cells were plunge-frozen into liquid ethane using a Leica EM GP2 automatic plunge-freezer. The ethane temperature was set to −183 °C and the chamber to 25 °C and 80% humidity. An additional 2 µl medium was added to the grid just before plunge-freezing. Grids were blotted from the back with Whatman® Type 1 paper for 3 s. Grids were clipped into FIB-AutoGrids™ (Thermo Fisher Scientific) designed for FIB milling.

**Cryo-light microscopy and cryo-focused ion beam milling**

Cryo-light microscopy was performed at -190°C using cryo-CLEM wide-field microscope (Leica Microsystems) equipped with a 50× objective with a numerical aperture of 0.9. A map was acquired as a Z-stack (30 µm, 300 nm spacing) in a bright field channel, blue fluorescence channel (excitation wavelength: 325-375 nm, emission wavelength: 435-490 nm) and green fluorescence channel (excitation wavelength: 450-490 nm, emission wavelength: 500-550 nm) covering 1.2 × 1.2 mm area using LAS X Navigator software (Leica). The final map was stitched using the Cryo-CLEM/Stitch TiltScan plugin available at https://github.com/Chlanda-Lab/cryoCLEM (*23*) in ImageJ/FIJI.

Cryo-focused ion beam milling was performed using Aquilos dual-beam cryo-focused ion beam-scanning electron microscope (cryo-FIB-SEM) (Thermo Fisher Scientific) with a cryo-stage cooled to -180 °C. Grids were mapped by cryo-scanning electron microscopy (cryo-SEM) and the cryo-LM map of the grid was correlated to the cryo-SEM map using the MAPS Software

(Thermo Fisher Scientific). Transfected cells were recognized by the mApple fluorescence signal and selected for milling. After the application of the protective organo-metallic platinum layer cells were milled gradually in 5 steps with a stage angle between 15° and 18° using a gallium ion beam. The first four steps were carried out automatically using a modified Autolamella script (*24*) available at https://github.com/Chlanda-Lab/autolamella. The last two milling steps were performed manually with a nominal thickness of 150 nm. Micro-expansion joints were used to minimize lamella bending (*25*).

**Cryo-electron tomography and tomogram reconstruction**

Cryo-electron tomography was done using a Krios cryo-TEM (Thermo Fisher Scientific) operated at 300 keV and equipped with a post-column Quantum Gatan Imaging energy filter (Gatan) and K3 direct electron detector (Gatan) with an energy slit set to 20 eV. As a first step, lamellae were mapped at 8,700× (pixel spacing of 10.64 Å) using a defocus of -65 µm in SerialEM (*22*) to localize double membrane vesicles. Tilt series were acquired using a dose-symmetric tilting scheme (*26*) with the zero-angle set to 8° and a nominal tilt range of 68° to -52° with 3° increments with SerialEM (*22*). Records were acquired as movies at target focus ranging from −4 to −2.5 µm, electron dose per record of 3 $e^{-}/Å^{2}$ and a magnification of 42,000× (pixel spacing of 2.156 Å). Beam-induced sample motion and drift were corrected using MotionCor2 (*27*). Tilt series was aligned using patch tracking or fiducials and tomograms were reconstructed using R-weighted back projection algorithm using 3DCTF, dose-weighting filter and SIRT-like filter 10 in the IMOD software package (*28*). Tomograms were denoised using Content Aware Image restoration (Cryo-CARE) (*29*). Figures show 3 or 5 slices of the tomogram averaged.

**Subtomogram averaging and tomogram rendering**

Pore densities were identified and extracted using a dipole model in Dynamo version 1.1.514 (*30*) using a box size of 256 pixels and an initial template model was created by averaging approximately 30 pores using the orientations inferred by the dipole model. To create the first average of the nsp3-4 sample, pores were picked using a general box model in Dynamo and aligned against the initial template using a spherical mask without imposing any symmetry. To create the average of the ΔUbl1-Mac1 and ΔUbl1-Ubl2 sample, a cylindrical mask was used.

The conical, azimuthal, translational search and angular increments were gradually decreased within 6 iterations using parameters for refinement of the average No.1 (Table S2). A symmetry scan was performed on the final average No.1 in Dynamo. Subsequently subtomogram averaging was performed using C6 symmetry where average No.1 was used as a new template to obtain an average No.2 using the same search parameters that were used for creating average No.1. The attained resolution was estimated using Fourier shell correlation with 0.5 and 0.143 criterion using a derived subtomogram averaging project with 2 references and from odd and even half-sets of particles. The C6-symmetrized map of average No.2 was visualized as isosurface in ChimeraX (*31*). The isosurface in Fig. 1 was low-pass filtered to 20Å whereas the isosurfaces in Fig. 4 and Fig. S6 were low-pass filtered to 40Å. The number of particles used for each nsp3-4 construct as well as the estimated resolution of all subtomogram averages is shown in Table S3. Volume rendering was performed manually in Amira (Thermo Fisher Scientific) after tomogram denoising using Cryo-CARE. Subtomogram average of the nsp3-4 pore was placed into the rendering based on coordinates determined from Dynamo cropping table using ArtiaX toolbox (*32*) in ChimeraX.

**Measurements and statistical analysis**

*Measurements of radius and curvature of DMVs.* The osculating circle at the interface of the DMVs were fitted manually in IMOD and the radius (R) of the osculating circle could be read directly. The mean curvature of the membrane was calculated as 1/R.

*Measurement of luminal spacing.* The luminal space of DMVs was measured as the distance between the outer leaflet of the outer membrane and the inner leaflet of the inner membrane. Line density profiles (~30 pixels in width, 0.41 Å/pixel) of ~30 nm in length were determined across the respective membrane using the plot profile tool in ImageJ/FIJI. The distances between the first and the fourth global maxima corresponding to the luminal space were then measured for each plot profile. For each DMV, three measurements were performed and the mean of them was reported.

*Measurement of the nearest distance between pores and number of pores per DMV.* The number of pores per DMV was counted manually in IMOD by creating models. Each project represented one tomogram, each contour represented a DMV, and each point represented a pore. The coordinates of each point were exported as .NFF files. The distance between each point and its nearest neighbor on the same contour was calculated by the script (Data S2).

Statistical analysis was done by using unpaired two-tailed t-test.

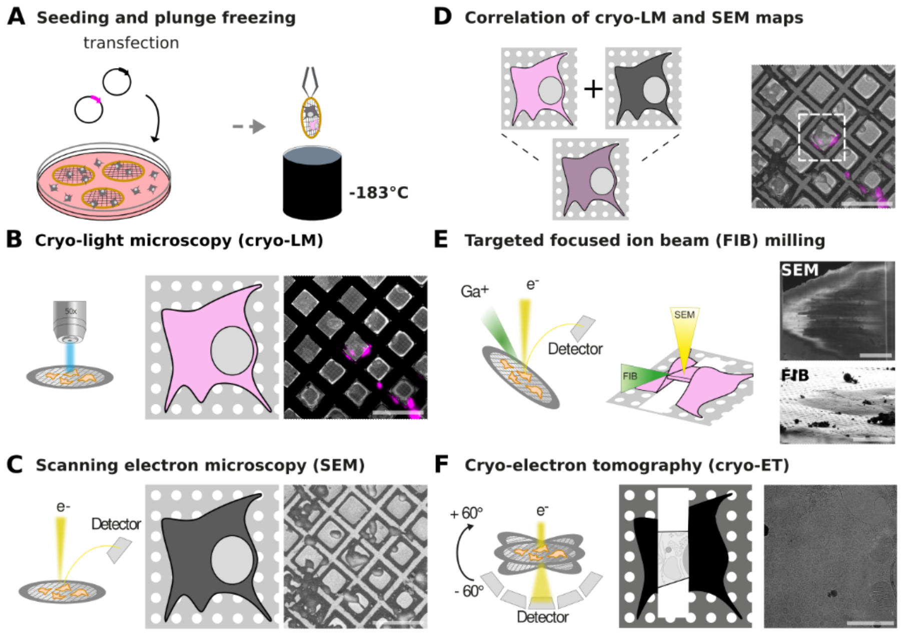


**Fig. S1.**
*In situ* cryo-correlative light and electron microscopy workflow for targeted cryo-focused FIB milling. (A) VeroE6 cells were seeded on gold EM grids and transfected with plasmids encoding for SARS-CoV-2 nsp3-4 proteins and a fluorescent marker (mApple) and plunge frozen in liquid ethane. (B) The grid was mapped in a cryo-LM. (C) Inside a cryo-FIB/SEM microscope, a high resolution SEM image was acquired. (D) The cryo-LM and SEM maps were correlated to find transfected cells with the fluorescent signal. (E) The cells of interest were FIB-milled. (F) The lamella was mapped by cryo-TEM and tilt series were acquired on sites of interest. Scale bars: (B-D) 200 µm; (E) SEM: 5 µm, FIB: 20 µm; (F) 1 µm. Edited from (*23*).

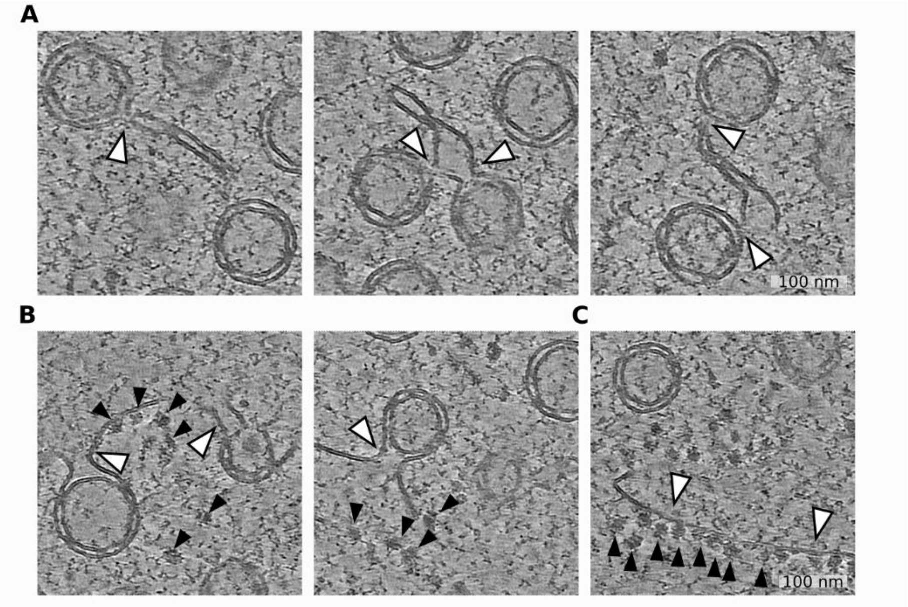


**Fig. S2.**
Gallery of DMCs, DMV-ER connections and ER cisternae with ribosomes in close proximity to DMV network. (A) Examples of DMCs containing pores interconnecting several DMVs indicated by white arrowheads. (B) DMVs connected to ER indicated by white arrowheads. Ribosomes are indicated by black arrowheads. (C) ER cisternae indicated by white arrowheads and ribosomes indicated by black arrowheads directly next to DMVs.

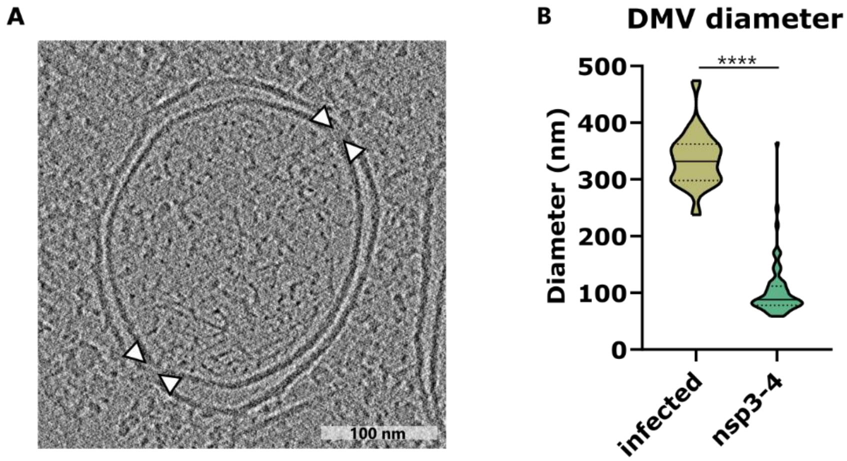


**Fig. S3.**
Comparison of DMV diameter in infected and nsp3-4 transfected cells. (A) Slice of a tomogram showing a DMV found in SARS-CoV-2 infected and chemically fixed VeroE6 cells (dataset of tomograms from a previous publication (*8*). Pores are highlighted by white arrowheads. (B) Plot showing diameter of DMVs in SARS-CoV-2 infected (n = 20, 16 hpi) and in nsp3-4 transfected cells (n = 62, 16 hpt). Data is shown as a Violin plot indicating the median, 25% and 75% quartiles. Unpaired two-tailed t-test showed significant differences with ****p<0.0001.

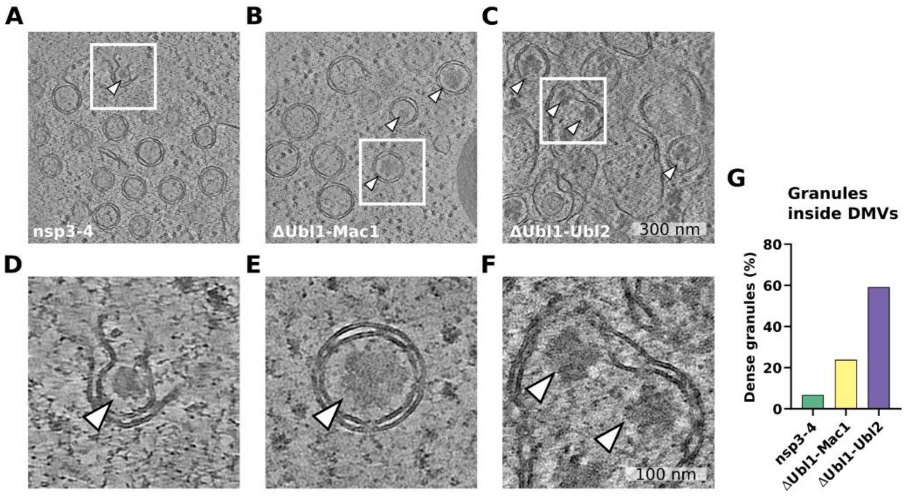


**Fig. S4.**
Granular density of unknown origin localized inside DMVs. (A-C) Averaged slices of tomograms of VeroE6 cells transfected with nsp3-4 (A), ΔUbl1-Mac1 (B) and ΔUbl1-Ubl2 (C) and processed at 18-24 hpt. (D-F) Averaged slices of tomograms showing the magnified views highlighted by a white rectangle in (A-C). Granular densities of unknown origin are indicated by white arrowheads. (G) Distribution of DMVs filled with granular densities compared to the total amount of DMVs in nsp3-4, ΔUbl1-Mac1 and ΔUbl1-Ubl2 samples.

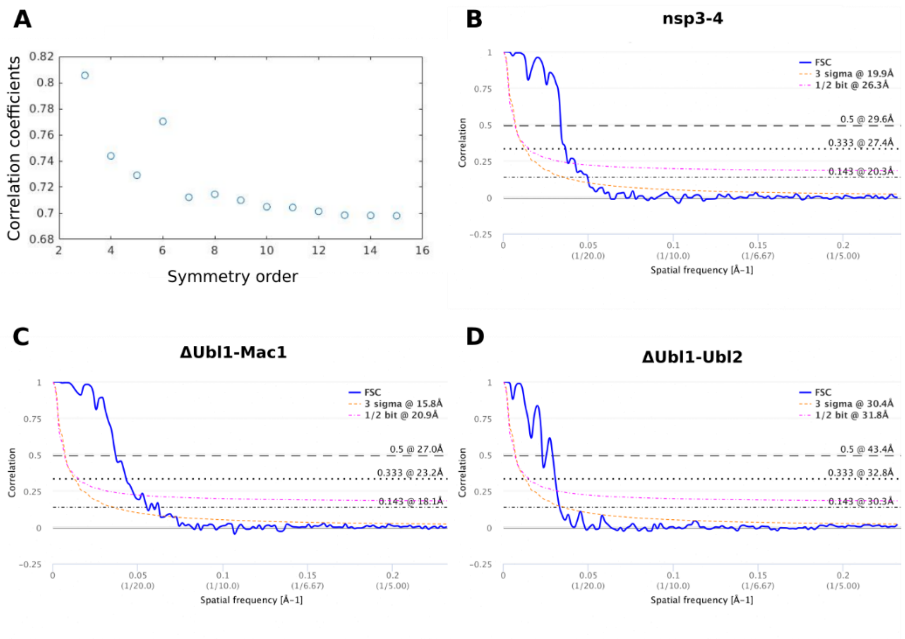


**Fig. S5.**
Fourier shell correlation curves from subtomogram averages upon implying sixfold symmetry. (A) Correlation coefficients between unsymmetrized and symmetrized averages. (B) Fourier shell correlation curve from nsp3-4 pores showing a resolution of around 20.3Å at 0.143 criterion. (C) Fourier shell correlation curve from ΔUbl1-Mac1 pores showing a resolution of around 18.1Å at 0.143 criterion. (D) Fourier shell correlation curve from ΔUbl1-Ubl2 pores showing a resolution of around 30.3Å at 0.143 criterion.

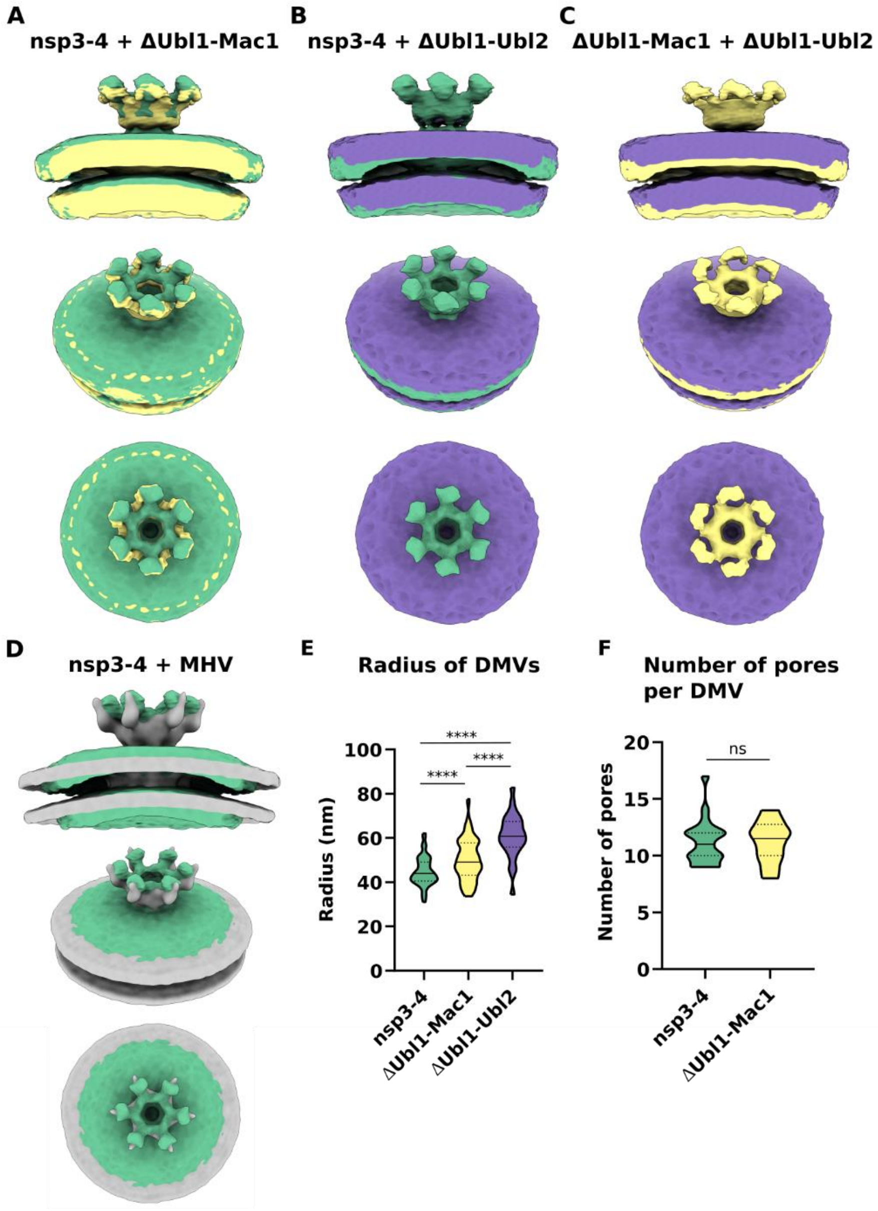


**Fig. S6.**
Comparison of isosurfaces and additional measurements characterizing the DMVs. (A) Isosurface of nsp3-4 pore compared to the isosurface of ΔUbl1-Mac1 pore showing partially collapsed prongs in ΔUbl1-Mac1 pore and differences in membrane curvature. (B) Isosurface of nsp3-4 pore compared to the isosurface of ΔUbl1-Ubl2 pore showing the loss of the prongs in ΔUbl1-Ubl2 pore and differences in membrane curvature. (C) Isosurface of ΔUbl1-Mac1 pore compared to the isosurface of ΔUbl1-Ubl2 pore showing the loss of the prongs in ΔUbl1-Ubl2 pore and differences in membrane curvature. (D) Isosurface of nsp3-4 pore compared to the isosurface of DMV pore of MHV infected cells showing thinner pore prongs induced by MHV and differences in membrane curvature. (E) Radius of DMVs in nsp3-4 (n = 103), ΔUbl1-Mac1 (n = 103), ΔUbl1-Ubl2 (n = 26). (F) Number of pores per DMV in nsp3-4 (n = 20), ΔUbl1-Mac1 (n = 20). Data is shown as a Violin plot indicating the median, 25% and 75% quartiles. Unpaired two-tailed t-test showed significant differences with ****$p<0.0001$.

| Step | T start | T end | Slope | Time | Reagent | Transfer | Agitation | UV |
|---|---|---|---|---|---|---|---|---|
| 1 | -90°C | -90°C | | 48 h | 0.1% UA | stay | off | off |
| 2 | -90°C | -45°C | 5 °C/h | 9 h | 0.1% UA | stay | off | off |
| 3 | -45°C | -45°C | | 5 h | 0.1% UA | stay | off | off |
| 4 | -45°C | -45°C | | 10 min | Acetone | exch/fill | off | off |
| 5 | -45°C | -45°C | | 10 min | Acetone | exch/fill | off | off |
| 6 | -45°C | -45°C | | 10 min | Acetone | exch/fill | off | off |
| 7 | -45°C | -45°C | | 4 h | 10% Lowicryl | mix | on | off |
| 8 | -45°C | -45°C | | 4 h | 25% Lowicryl | mix | on | off |
| 9 | -45°C | -35°C | 2.5 °C/h | 4 h | 50% Lowicryl | mix | on | off |
| 10 | -35°C | -35°C | 2.5 °C/h | 4 h | 75% Lowicryl | mix | on | off |
| 11 | -35°C | -35°C | | 10 h | 100% Lowicryl | exch/fill | off | off |
| 12 | -35°C | -35°C | | 10 h | 100% Lowicryl | exch/fill | off | off |
| 13 | -35°C | -35°C | | 10 h | 100% Lowicryl | exch/fill | off | off |
| 14 | -35°C | -35°C | | 48 h | 100% Lowicryl | stay | off | on |
| 15 | -35°C | 20°C | 5 | 9 h | 100% Lowicryl | stay | off | on |
| 16 | 20°C | 20°C | | 48 h | 100% Lowicryl | stay | off | on |
| 17 | 20°C | 20°C | | 72 h | 100% Lowicryl | stay | off | off |

**Table S1.**
Automated freeze substitution program. UA: uranyl acetate.

| Parameters | Round 1 | Round 2 | Round 3 | Round 4 | Round 5 | Round 6 |
|---|---|---|---|---|---|---|
| Iterations | 1 | 1 | 1 | 1 | 1 | 1 |
| References | 1 | 1 | 1 | 1 | 1 | 1 |
| Cone aperture [°] | 360 | 180 | 90 | 45 | 24 | 12 |
| Cone Sampling [°] | 120 | 60 | 30 | 15 | 8 | 4 |
| Azymuth rotation range [°] | 360 | 180 | 90 | 45 | 24 | 12 |
| Azymuth rotation sampling [°] | 120 | 60 | 30 | 15 | 8 | 4 |
| Refine | 5 | 5 | 5 | 5 | 5 | 5 |
| Refine factor | 2 | 2 | 2 | 2 | 2 | 2 |
| High pass filter [pixel] | 2 | 2 | 2 | 2 | 2 | 2 |
| Low pass filter [pixel] | 32 | 32 | 64 | 64 | 80 | 128 |
| Symmetry | C1/C6 | C1 / C6 | C1 / C6 | C1 / C6 | C1 / C6 | C1 / C6 |
| Particle dimensions | 128 | 128 | 128 | 128 | 128 | 256 |
| Shift limits in X, Y and Z [pixel] | 30, 30, 30 | 20, 20, 20 | 12, 12, 12 | 8, 8, 8 | 4, 4, 4 | 2, 2, 2 |
| Shift limiting way | 2 | 2 | 2 | 2 | 2 | 2 |
| Separation in tomogram | 1 | 1 | 1 | 1 | 1 | 1 |

**Table S2.**
Numerical parameters for subtomogram averaging calculation in Dynamo.

| Construct | Number of tomograms | Number of particles | Resolution (0.143 Criterion) | Resolution (0.5 Criterion) |
|---|---|---|---|---|
| Nsp3-4 | 8 | 325 | 20.3 Å | 29.6 Å |
| Nsp3-4 ΔUB1-MAC1 | 7 | 1099 | 18.1 Å | 27.0 Å |
| Nsp3-4 ΔUB1-UBl2 | 8 | 94 | 30.3 Å | 43.4 Å |

**Table S3.**

Number of particles used for subtomogram averaging and attained resolution.

**Movie S1.**

Cryo-electron tomogram and rendering of DMVs formed in VeroE6 cells expressing nsp3-4 construct.

**Movie S2.**

Cryo-electron tomogram of DMVs formed in VeroE6 cells expressing ΔUbl1-Mac1 construct.

**Movie S3.**

Cryo-electron tomogram and rendering of DMVs formed in VeroE6 cells expressing ΔUbl1-Ubl2 construct.

**Movie S4.**

Cryo-electron tomogram and rendering of DMVs formed in VeroE6 cells expressing GG>AA construct.

**Data S1. (separate file)**
Protein sequences of nsp3-4 constructs

**Data S2. (separate file)**
Script for calculation of pore-to-pore nearest distance
